# Altering stimulus timing via fast rhythmic sensory stimulation induces STDP-like recall performance in human episodic memory

**DOI:** 10.1101/2022.11.02.514843

**Authors:** Danying Wang, Kimron L. Shapiro, Simon Hanslmayr

## Abstract

Animal studies suggest that the strength of synaptic modification depends on spike timing between pre- and post-synaptic neurons on the order of tens of milliseconds, which is termed ‘spike-timing-dependent plasticity’ (STDP). However, evidence for STDP in human episodic memory is lacking. We investigated this using rhythmic sensory stimulation to drive visual and auditory cortices at 37.5 Hz with four phase offsets. Visual relative to auditory cued recall accuracy was significantly enhanced in the 90° condition since the visual stimulus led at the shortest delay (6.67 ms). This pattern was reversed in the 270° condition when the auditory stimulus led the shortest delay. Within cue modality, recall was enhanced when a stimulus of the corresponding modality led the shortest delay as compared to the longest delay (20 ms). Our findings provide novel evidence for STDP in human memory, which builds an important bridge from in-vitro studies in animals to human behaviour.

## Introduction

Episodic memory provides humans with the ability to mentally travel back to the past (Tulving, 2002) where experiences typically involve associations between multimodal information. For example, when you stopped by a beautiful mural painted on a wall last Sunday, you heard jazz music being played from a pub on the other side of the street. Forming a memory of the association between those elements is thought to depend on modification of synaptic connectivity (Martin et al., 2000; Neves et al., 2008). Laboratory animal studies provide correlational and causal evidence supporting the link between synaptic plasticity and the formation of long-term memory (Bliss & Lømo, 1973; Nabavi et al., 2014; Whitlock et al., 2006). However, and to the point of the present study, much less is known about if and how similar mechanisms play a role in human episodic memory.

Donald Hebb proposed synaptic plasticity as the neuronal basis for learning and memory in his seminal book (Hebb, 2002), writing ‘Neurons that fire together wire together’. In line with this postulate, a form of synaptic plasticity that requires two neurons firing within a time window on the order of tens of milliseconds to induce a synaptic change has been found (Bi & Poo, 1998; Markram et al., 1997). This form of synaptic plasticity suggests that the temporal interval and order between pre-synaptic and post-synaptic spikes are the determining factors for the direction of synaptic modification, which is termed spike-timing dependent plasticity (STDP) (Caporale & Dan, 2008). Specifically, long-term potentiation (LTP) can be induced if the pre-synaptic neuron fires within a critical time window before the post-synaptic neuron. On the other hand, reversing the order weakens the synaptic strength, leading to long-term depression (LTD).

Evidence for STDP has come pre-dominantly from animal studies *in-vitro* and *in-vivo* (Caporale & Dan, 2008), with far fewer studies showing evidence for STDP in the human brain (Mansvelder et al., 2019). Human in-vitro studies conducted on brain tissue show that the strength of synaptic modification depends on the relative timing between pre-synaptic and post-synaptic activity albeit with different time scales, as compared to the critical time window of STDP in animals (Testa-Silva, 2010; Verhoog et al., 2013), which begs the question to what extent the findings in animals apply to the healthy human brain. STDP-like effect has been found in the human motor cortex (Stefan, 2000; Wolters et al., 2003), somatosensory cortex (Wolters et al., 2005), as well as in frontal networks (Guidali et al., 2021), using a transcranial magnetic stimulation (TMS) protocol called paired associative stimulation (PAS) (Müller-Dahlhaus, 2010). Additionally, human psychophysics experiments suggest that precisely controlling the relative timing of two visual inputs within a time window of 40 ms, results in a perceptual pattern resembling STDP (Fu et al., 2002; Yao et al., 2004; Yao & Dan, 2001).

Whereas the above studies suggest that STDP is present in the human brain, strong evidence about its involvement in the formation of episodic memory is missing. This is somewhat surprising given Hebb’s influential theory and the dominance of STDP in animal models of memory. A recent study using a rhythmic sensory entrainment approach (Hanslmayr et al., 2019) precisely controlled the input timing relative to ongoing theta phase and provides direct evidence on the role of theta-phase mediated plasticity in human episodic memory (Clouter et al., 2017). Nevertheless, it is not known if human episodic memory performance is influenced by relative timing between inputs on the order of tens of milliseconds, which is the temporal characteristic that determines STDP. In the present study, we examined this question using rhythmic sensory entrainment at 37.5 Hz, which allowed us to alter the relative timing between visual and auditory stimuli at fine temporal resolution with four phase offset conditions, 0°, 90°, 180° and 270° (Fig. 1b). We tested participants’ cued recall accuracy on the association between visual and auditory stimuli to investigate the role of STDP on bi-directional association.

**Figure 1.**
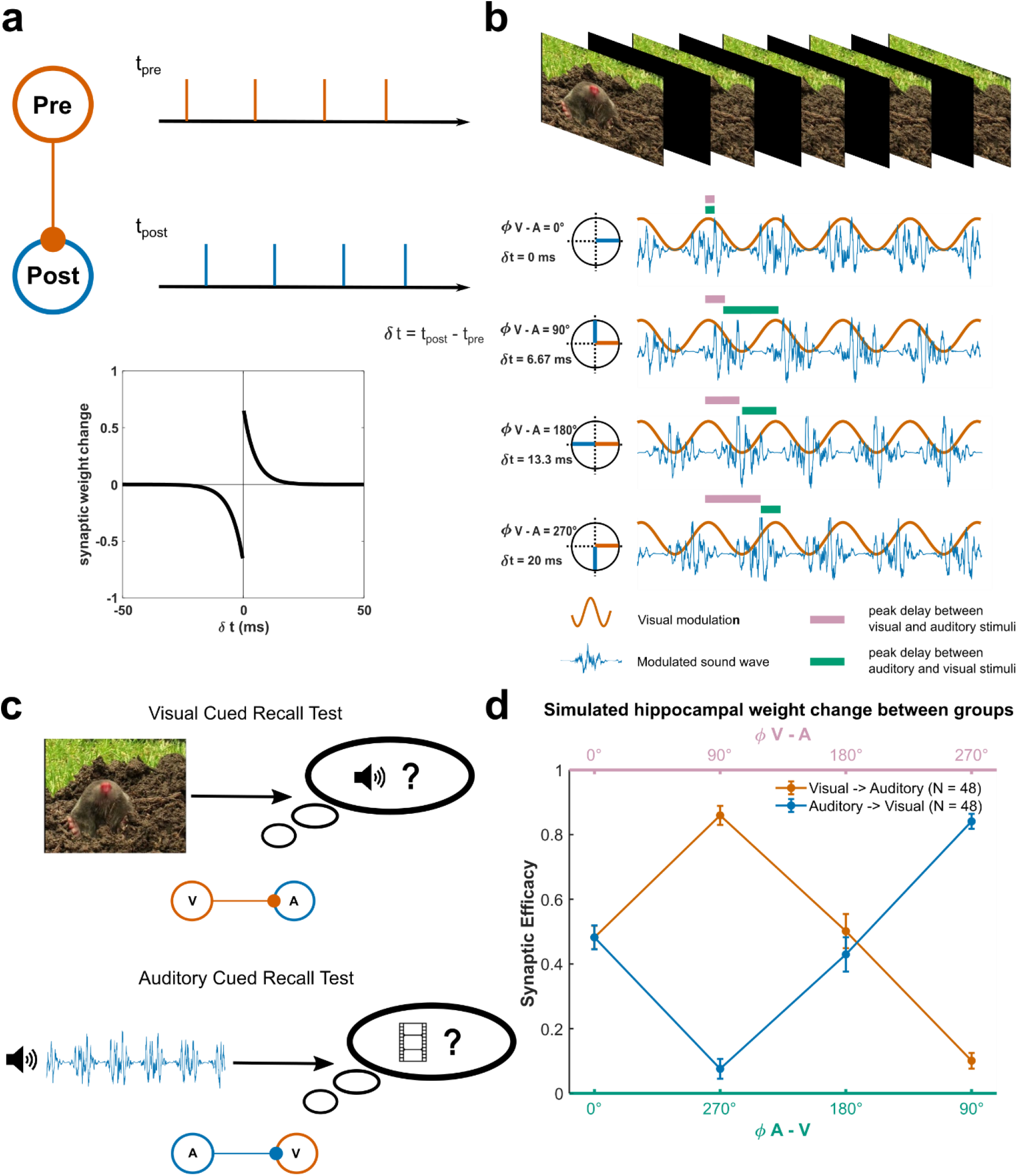
Experiment design. **(a)** Schematic of the STDP framework. Synaptic weight change depends on spike timing between a pre-synaptic neuron and a post-synaptic neuron. LTP occurs if the post-synaptic neuron fires in a short delay after pre-synaptic neuron firing, while LTD happens if the order of firing is reversed. **(b)** The encoding phase involved viewing a 3 s video and listening to a 3 s sound clip. The luminance and amplitude of videos and sounds were modulated at 37.5 Hz with four phase offsets, 0°, 90°, 180°, and 270°. The pink bars represent peak delays between a video and a sound, which were 0, 6.67 ms, 13.33 ms and 20 ms for 0°, 90°, 180°, and 270° offset conditions, respectively. The teal bars represent peak delays between a sound and a video, which were 0, 20 ms, 13.33 ms and 6.67 ms, respectively. **(c)** Visually cued recall presented the video in the memory test phase. Participants were asked to select the correct sound. In the auditorily cued recall experiment, participants were cued with the sound in the test phase and asked to recall the paired video. **(d)** Hippocampal weight changes between two groups of neurons as a function of phase offset conditions, simulated by a computational model that implemented the STDP learning rule. The input stimuli were modulated by a 37.5 Hz sine wave at the same phase offset conditions as in the experiments. The weights were averaged across 48 simulations for each condition. Error bars represent standard error (SE).

## Results

We stimulated auditory and visual cortices at 37.5 Hz with four phase offsets between the two sensory stimuli, 0°, 90°, 180° and 270° (Fig. 1b). Accordingly, the visual inputs precede the auditory inputs by 0 ms, 6.67 ms, 13.33 ms or 20 ms, respectively. The corollary auditory inputs precede the visual inputs by 0 ms, 20 ms, 13.33 ms or 6.67 ms, respectively. The auditory and visual stimuli were music clips and documentary video clips, respectively, that were not obviously related to each other. Participants were asked to memorise the pairs of video and sound clips. During the recall phase, we cued participants’ memory with each stimulus modality in a between design. In group 1, participants were cued with a visual stimulus and asked to recall the paired auditory stimulus. Participants in group 2 were cued with an auditory stimulus and asked to recall the paired visual stimulus (Fig. 1c). Our main prediction was that recall accuracy decreases or increases as a function of (i) phase offset between auditory and visual stimuli and (ii) the modality of the memory cue.

### Simulating the results with an STDP computational model

To formalise the predictions of STDP for our experimental results, we first simulated our experimental paradigm with a computational model that implements the STDP learning rule. Two groups of neurons that are simulated by an integrate-and-fire equation (cf. Parish et al., 2018; Wang et al., 2021) received two stimuli. The stimuli were modulated at 37.5 Hz with four phase offsets: 0°, 90°, 180°, and 270°, which corresponded to the visual and auditory stimuli in our experiments. Synaptic weights from one group of neurons (e.g., visual) to the other group of neurons (e.g., auditory) were rewarded if visual neurons fired before auditory neurons and punished if auditory neurons fired first. The weight changes decayed exponentially over time (Song et al., 2000).

To evaluate model recall performance, weights from the visual group to the auditory group, and from the auditory group to the visual group, were averaged across 48 simulations for each phase offset condition. The model reveals that the weights from the visual group to the auditory group were higher compared to the weights from the auditory group to the visual group when the visual stimulus led the auditory stimulus by 90° (6.67 ms). This pattern was reversed when the visual stimulus led the auditory stimulus by 270° (20 ms), which means that the auditory stimulus led the visual stimulus by 90° (Fig. 1d). The weights did not differ between 0° and 180° phase offset conditions, which might appear counterintuitive for the 0° condition. The outcome of this particular condition is caused by the fact that half the pre-synaptic spikes precede the post-synaptic spikes while half the pre-synaptic spikes follow the post-synaptic spikes, thus causing reward and punishment on weights to cancel each other. Consistent with this model we therefore predicted the strongest memory differences to occur between the 90° and 270° conditions, depending on the modality of the cue and the target (i.e., to-be-recalled) memory. Specifically, visually cued recall accuracy should be higher than auditorily cued recall accuracy when the visual stimulus led the auditory stimulus by 90°. This pattern should be reversed in the condition where the visual stimulus led the auditory stimulus by 270° (i.e., when the auditory stimulus led the visual stimulus by 90°). Moreover, visually cued recall accuracy should be better when the visual stimulus led the auditory stimulus by 90° compared to 270°, while the pattern should be reversed for auditorily cued recall accuracy. No difference was expected between visually cued recall accuracy and auditorily cued recall accuracy in the 0° and 180° conditions.

### Recall accuracy as a function of the actual phase difference between entrained visual and auditory gamma activity

EEG was recorded for 24 participants in each group, which allowed us to confirm that visual and auditory brain activity was modulated by the sensory stimuli at 37.5 Hz at the specified phase offsets. It is important to note the difference in transduction delays from the eye/ear to the brain, whereby the auditory domain reaches the cortex before the visual domain by approximately 40 ms (Di Russo et al., 2002; Picton, 2010; Schnapf et al., 1987). Therefore, we added a 40 ms delay before the auditory stimulus onset to approximate simultaneous processing in visual and auditory brain regions (Clouter et al., 2017; Wang et al., 2021). However, given that the modulation frequency of 37.5 Hz is relatively fast, just a few milliseconds difference in the transduction delays would cause the phase modulation to fail (for example, 5 ms corresponds to 67.6° at 37.5 Hz).

To compensate for the above problem, we first computed the phase differences between visual and auditory regions in each experimental condition to label each condition according to its actual phase offset. In the second step, source activity was reconstructed from each ROI (Fig. 2a), and the ERP was computed for each condition and each ROI. Next, the grand averaged ERP was computed for all participants, given that stimulus modulation in the encoding phase was the same for both groups. Fig. 2b reveals that the actual instantaneous phase differences between visual and auditory grand average ERPs were 180° off from the expected phase offset conditions. The experimental conditions 180°, 270°, 0° and 90° showed 0°, 90°, 180°, and 270° phase differences between visual and auditory activity, respectively. This observation was confirmed statistically by both Rayleigh and V tests. In these tests 1025 data points (0.5 and 2.5 s) of instantaneous phase difference between auditory and visual signals were used as the dependent variable. The null hypothesis of uniform distribution was rejected for all conditions (all *p* values ≈ 0). Moreover, the mean direction was confirmed to be 0°, 90°, 180°, and 270° in the experimental conditions 180°, 270°, 0° and 90°, respectively (V test; all *p* values = 0).

**Figure 2.**
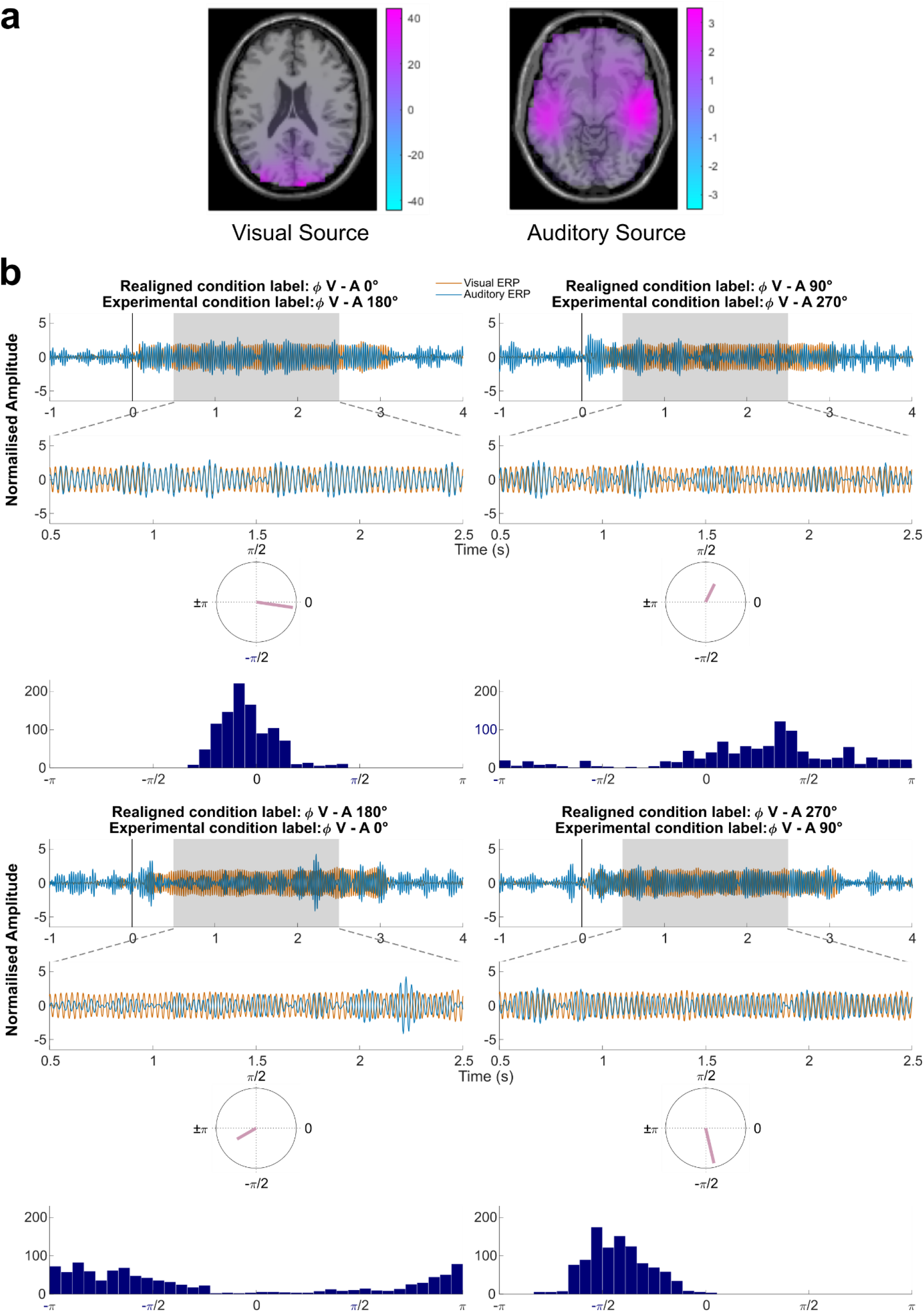
Phase differences between visual and auditory grand average ERP in each phase offset condition. **(a)** Source localisation of visual and auditory sources in the unimodal conditions. Visual source, MNI coordinates of regions-of-interest (ROI): 10, -99, 20. Auditory sources, MNI coordinates of ROIs: right, 50, -19, -10; left, -50, -31, 0. Unimodal stimuli were modulated at 4 Hz. Evoked power was averaged between 3.5 and 4.5 Hz, and between 0.75 and 2.75 s at each virtual electrode in the unimodal visual and auditory conditions. The values were normalised by averaged evoked power of pseudo baseline conditions which the trials were selected randomly to shift by 0°, 90°, 180° or 270° (see Methods). **(b)** Phase differences between visual and auditory sources in each phase offset condition. Grand average ERP signals (N = 48) at visual and auditory sources were band-pass filtered at 35 to 40 Hz. Amplitude was normalised. Instantaneous phase differences were averaged between 0.5 and 2.5 s (shaded time window) and plotted by the mean resultant vector on a unit circle. Histograms showed wrapped instantaneous phase differences between 0.5 and 2.5 s.

Given the outcome of these tests, we relabelled our experimental conditions based on the actual phase offsets before investigating recall accuracy. Recall accuracy in each condition for each participant was normalised by their mean performance. Fig. 3a shows the mean normalised recall accuracy of 48 participants in each group for each newly labelled condition. We consistently found that the visually cued recall accuracy in the actual 90° phase offset condition was higher than the auditorily cued recall accuracy. An independent samples t-test confirmed this difference to be statistically significant, *t*(94) = 2.065, *p* = 0.021 (one-tailed), Cohen’s *d* = 0.421. An independent samples t-test showed that in the 270° phase offset condition, the auditorily cued recall accuracy did not differ significantly from the visually cued recall accuracy (*t*(94) = -0.1622, *p* =0.436 (one-tailed)). Further, we analysed participants’ subjective rating on how well a given sound suited the content of the corresponding video. A main effect of subsequent memory revealed that the mean rating for subsequently remembered pairs was significantly higher than for subsequently forgotten pairs, *F*(1, 94) = 47.113, *p* < 0.001, reflecting that video-sound pairs that were perceived as congruent were remembered better. However, the rating did not differ between the visually cued group and the auditorily cued group for the 90° phase offset condition, which suggests the memory effect observed in the 90° phase offset condition was not driven by higher semantic congruency between video-sound pairs in the visually cued recall group. In summary, the results in the large sample (N = 96, with N = 48 per cue condition) resemble a ‘glass half full or half empty’ outcome, with the 90° phase offset condition being consistent with the model predictions but the 270° phase offset condition being inconsistent with those predictions.

**Figure 3.**
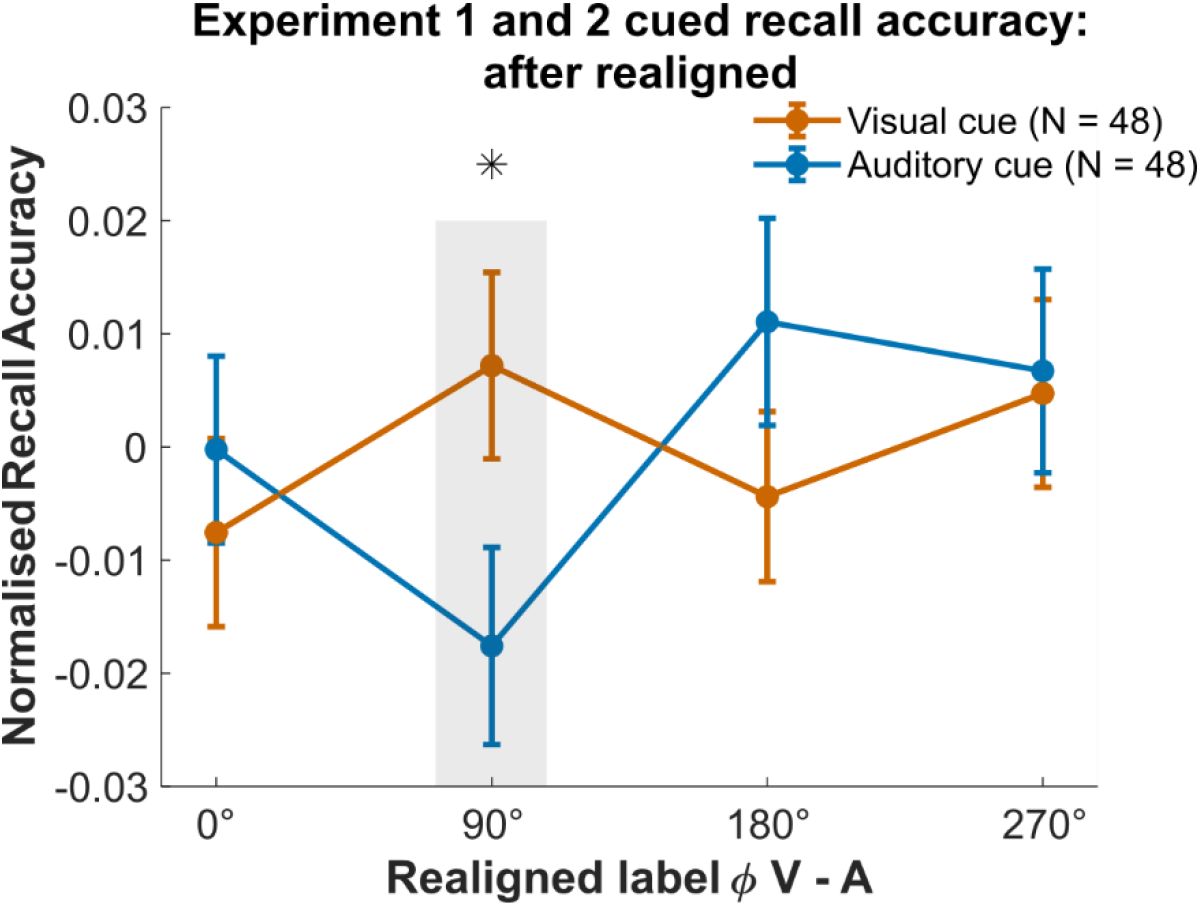
Behavioural results based on the actual phase difference between visual and auditory grand average ERP. Normalised recall accuracy in each phase offset condition that showed the actual phase differences 0°, 90°, 180°, and 270° between visual and auditory grand average ERP, respectively. Error bars represent SE. * *p* < 0.05. N = 48 for both groups.

### Recall accuracy as a function of phase difference between single trial visual and auditory gamma activity

We investigated the relationship between stimulus timing and recall behaviour on the single trial level. Notably, Wang et al. (2018) observed considerable trial-by-trial variation of phase differences arising between auditory and visual cortices even though the phase difference of the sensory stimulation was constant. Therefore, we investigated if such a trial-by-trial variance in phase differences between entrained visual and auditory source activity contributed to the discrepancy between the behavioural results and the model predictions. Only participants with EEG recordings were used for this analysis (N=48; with N=24 per cue condition). Instantaneous phase differences between band-pass filtered (35 and 40 Hz) visual and auditory sources were averaged between 0.5 and 2.5 s for each trial. Based on this phase difference, single trials were sorted between -π and π and divided into four equally sized bins (Fig. S2). This procedure was implemented to realign each trial to the actual onset of sensory responses to the external stimuli, thus reducing the influences of any factors that could cause a variation of phase differences (or transduction delays). As predicted, after sorting the trials into four phase bins, the resulting grand average ERP in each phase bin showed significant phase concentration at the intended phase directions, as confirmed by V test (Fig. 4a). Both Rayleigh test and V test on 1025 data points in each phase bin rejected the null hypothesis of uniform distribution. All p values for the statistical tests were ≈ 0.

**Figure 4.**
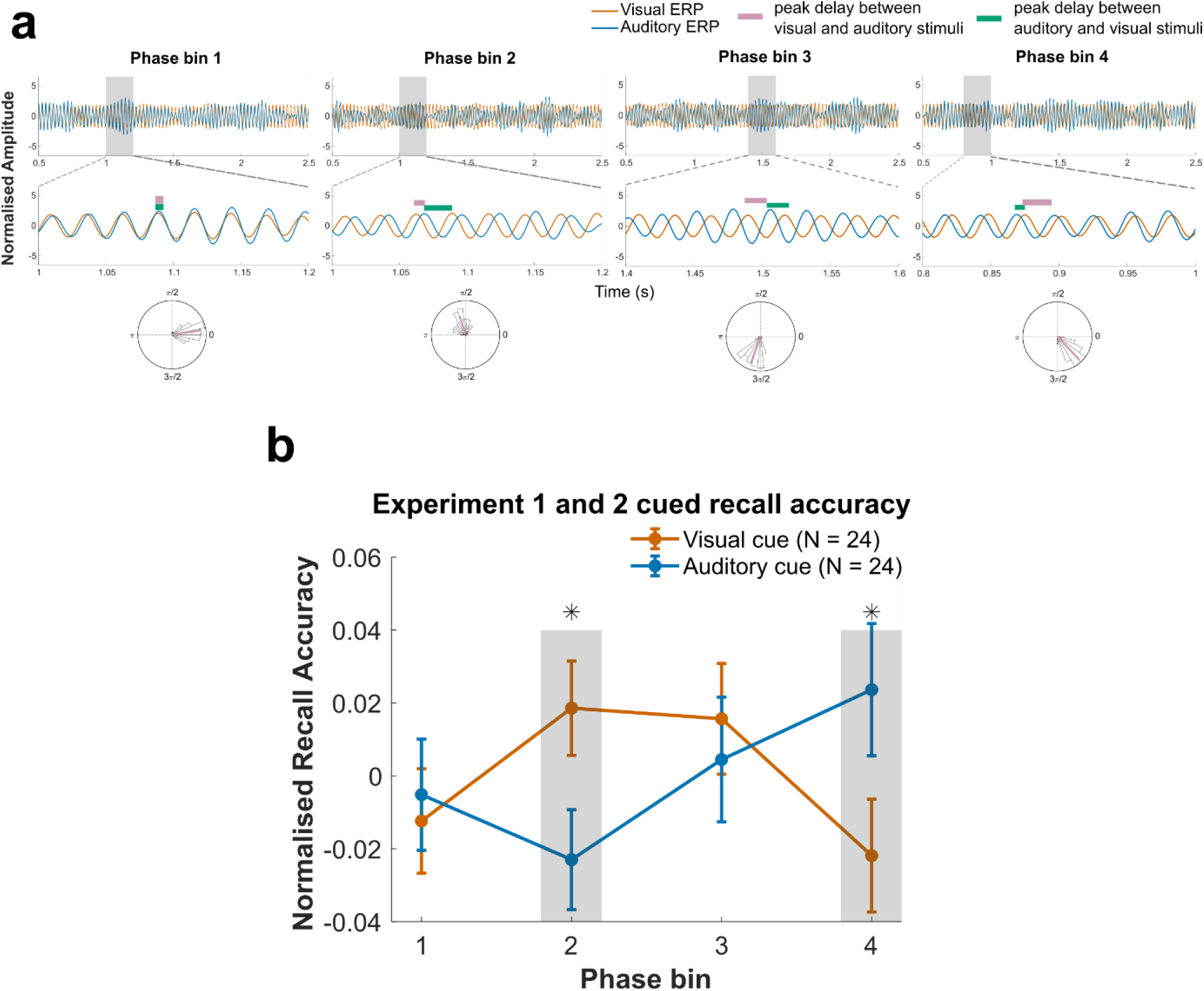
Recall accuracy as function of single trial phase offset between visual and auditory activity. **(a)** Phase differences between visual and auditory grand average ERP (N=48) in each phase bin. Amplitude was normalised. Instantaneous phase differences were averaged between 0.5 and 2.5 s and plotted by the mean resultant vector on a unit circle with count histogram of the phase differences between 0.5 and 2.5 s. The selected (grey shaded) time windows were zoomed in to show the phase delays between visual and auditory time series. **(b)** Normalised recall accuracy in each phase bin. Error bars represent SE. * p < 0.05. N = 24 for both groups.

After single-trial sorting, the proportion of remembered trials in each phase bin was calculated and normalised as above. Fig. 4b shows the mean normalised recall scores in each phase bin for each group of participants. Importantly, the resulting pattern resembles the simulated data from the STDP model with the normalised recall score in phase bin 2 (i.e., 90°) higher for the visually cued group compared to the auditorily cued group. In contrast, this pattern was reversed in phase bin 4 (i.e., 270°) where the auditorily cued group revealed better memory compared to the visually cued group. An ANOVA with the repeated measures factor ‘phase bin’ (phase bin 2 vs. phase bin 4) and a between subject factor ‘cue condition’ (visual vs. auditory) indicated a significant interaction, *F*(1,46) = 6.444, *p* = 0.015, η^2^_*p*_ = 0.123. Independent samples t-tests confirmed statistically that the normalised recall score in the visually cued group was significantly better than in the auditorily cued group in phase bin 2, *t*(46) = 2.203, *p* = 0.016 (one-tailed), Cohen’s *d* = 0.636, whereas the pattern was reversed in phase bin 4, *t*(46) = -1.908, *p* = 0.024 (one-tailed), Cohen’s *d* = -0.551. Paired samples t-tests confirmed the hypothesis that visually cued recall is better when the visual stimulus led by the shortest delay, i.e., 90° or 6.67 ms relative to when the visual stimulus led by the longest delay, i.e., 270° or 20 ms. The pattern was predicted to be reversed for auditory recall relative to when the visual stimulus led by 90° or 270°, i.e., the auditory stimulus should lead by 270° or 90°, respectively. Indeed, recall accuracy in phase bin 2 was significantly higher than in phase bin 4 in the visually cued condition, *t*(23) = 1.877, *p* = 0.037 (one-tailed), Cohen’s *d* = 0.383. This pattern was completely reversed in the auditorily cued condition where recall accuracy in phase bin 4 was higher compared to phase bin 2, *t*(23) = -1.747, *p* = 0.047 (one-tailed), Cohen’s *d* = -0.357. Moreover, the analysis of participants’ subjective rating on how well a sound suited the contents of a video suggests that the rating scores did not differ between recall cued groups in either phase bin 2 or phase bin 4, nor between phase bins in each group. Neither was a significant interaction found between phase bin and cue condition. Together, these results suggest that trial-by-trial phase differences between rhythmically stimulated visual and auditory sources generate memories that are consistent with the results simulated by a computational model implementing a STDP learning rule.

## Discussion

Forming lasting associations of stimuli on a one-shot basis, which is the hallmark of episodic memory, is thought to depend on synaptic plasticity. Spike-timing dependent plasticity is one of the best documented plasticity mechanisms in animals and emphasizes that the temporal order and interval between two spike events determine the direction and strength of synaptic modification. Our results are the first to reveal that stimulus timing on the order of milliseconds influences human episodic memory in a way that is highly consistent with STDP. This was achieved by modulating visual and auditory stimuli at 37.5 Hz with 0°, 90°, 180° and 270° phase offsets. The mean memory score for trials that showed visual brain activity leading auditory activity by 90° (∼6.67 ms) was significantly higher when memory was cued with a visual stimulus, than when memory was cued with an auditory stimulus. The pattern was reversed when memory score was computed for the trials where auditory activity led the visual activity by 90°. Importantly, our findings are consistent with simulations from a computational model of STDP, in turn suggesting that STDP can account for the observed reversal of memory performance when cued with different modalities. Together, the results provide evidence that precise timing and order between sensory inputs is crucial for encoding multisensory information into episodic memory, which provides strong evidence for a role of STDP in human episodic memory.

Previous studies show that human synapses also follow the STDP rule, although the time window for inducing LTP spans across negative and positive intervals, which is wider than the classic STDP rule observed in juvenile rats (Bi & Poo, 1998; Caporale & Dan, 2008; Mansvelder et al., 2019; Testa-Silva, 2010; Verhoog et al., 2013). These *in-vitro* studies isolated brain slices from epilepsy or tumour patients with wide age ranges. It is still unclear whether synapses in young and intact human brains show STDP properties similar to rodents. In the current study, we simulated our experimental procedure with a simple spiking model that implements the classic STDP learning rule (Parish et al., 2018; Wang et al., 2021). In our model, synaptic weights are strengthened if a spike of the pre-synaptic neuron arrives before a spike of the connected post-synaptic neuron, whereas the weights are weakened when this sequence is reversed. Sorting recall accuracy into four phase bins from single trials resembled the pattern of the synaptic weight changes in each phase offset condition simulated by our model, which suggests that the classic STDP learning rule is sufficient to account for the memory results of the current study. Human *in-vivo* studies have shown that stimulus timing and order modulate behavioural changes in perception using PAS and paired sensory stimulation (Fu et al., 2002; Romei et al., 2016; Yao et al., 2004; Yao & Dan, 2001). The direction and degree of the behavioural changes resemble stimulus-timing dependent cortical changes found *in-vivo* in animal brains (Fu et al., 2002; Yao et al., 2004; Yao & Dan, 2001; Zhang et al., 1998). Though the exact time windows where the largest changes are induced differ depending on the cortical regions, the time windows when the changes are exerted fall within the STDP time frame. Notably, perceptual enhancement has always been found when the presentation of stimuli is near-synchronous rather than synchronous, which is indeed consistent with physiological data (Bi & Poo, 1998; Fu et al., 2002; Yao et al., 2004; Yao & Dan, 2001). Our results are the first to demonstrate that near-synchronous cross-modality stimulus presentation enhances or impairs associations in episodic memory depending on which modality was leading. The association from visual to auditory stimuli was enhanced when the visual stimulus led the auditory stimulus by 6.67 ms, whereas the association from auditory to visual stimulus was impaired.

Using the same memory paradigm, Clouter et al. (2017) and Wang et al. (2018) showed that synchronous stimulus presentation enhances episodic memory formation. However, the temporal resolution in those studies is considerably lower than in the current study as stimulus inputs were modulated at theta frequency (4 Hz). We speculate that spike timing might be coordinated with theta oscillations, thus interacting with the theta-phase dependent learning mechanism (Cobb et al., 1995; Huerta & Lisman, 1995; Wang et al., 2021). The synchronous condition in the studies from Clouter et al. (2017) and Wang et al. (2018) allows firing at the LTP-inducing theta phase and the individual pairs of spike timing fall within the STDP window. Indeed, memory performance was significantly worse for those conditions where the peak delays were outside of the STDP window, i.e., >= 90° phase difference (62.5 ms) compared to the synchronous condition. Using computational modelling, Wang et al. (2021) also demonstrate that STDP is involved in the synchronous theta-phase memory effect. However, given that memory was cued only uni-directionally in Clouter et al. (2017) and Wang et al. (2018), we could not experimentally disentangle the role of STDP due to the low temporal resolution, which in turn underscores the importance of the current study.

Our results suggest that the preferred direction of association can be determined by stimulus presentation order. The preference of forward presentation order for learning and associative memory has been found on a large time scale, in the order of seconds (Kahana, 1996; Kahana & Caplan, 2002; Tanimoto et al., 2004). However, STDP suggests that association happens on a much more precise time scale. Therefore, the direct link between those findings on learning and associative memory and STDP is still not clear (but see Drew & Abbott, 2006; Reifenstein et al., 2021; Skaggs et al., 1996 for possible links between STDP and learning that happen in sequences on a larger time scale). The difference in peak delay between the 90° phase offset condition and the 270° phase offset condition in the current study is in the order of tens of milliseconds (6.67 ms vs. 20 ms), which reveals that the preferred direction of association can be induced by STDP. Moreover, methodologically we provide the novel insight that human episodic memory formation can be manipulated by stimulus timing on a fast time scale that is within the STDP window.

Previous electrophysiology and human psychophysics studies suggest that stimulus-timing induced neuronal changes directly result in changes in perception (Caporale & Dan, 2008). We did not directly measure neuronal changes such as tuning of neurons or synaptic modification in the current study; however, simulated results from our model suggest that synaptic weights between the two groups of neurons change as a function of stimulus phase offset condition following learning. Our behavioural results based on the same stimulus inputs are consistent with the simulated synaptic weight changes. Interestingly, we found a modality preference for the presentation order and interval at corresponding sensory cortices, suggested by the inter-trial-phase-coherence (ITPC) at the stimulation frequency 37.5 Hz (Fig. S1). In the visual cortex, ITPC in the 90° phase offset condition in which the visual stimulus led, was significantly stronger than in the 270° phase offset condition. Whereas in the auditory cortex, ITPC in the 90° phase offset condition was significantly weaker than in the 270° phase offset condition. The results are in line with previous findings on cortical STDP, suggesting a directionality specific excitability in the corresponding sensory cortices (Lakatos et al., 2009; Müller-Dahlhaus, 2010). However, it cannot explain the memory effect since the modality preference was consistent across visual and auditory cue groups because of lack of interaction between brain region, phase offset condition and cue group (Fig. S1). To the extent that the modality preference effect can be linked to memory requires further investigation. Given that our experimental task is highly hippocampus-dependent, requiring the binding of two semantically unrelated stimuli of two modalities into long-term memory in a single trial, the results we find are consistent with STDP in the hippocampus (Bi & Poo, 1998).

A recent intracranial EEG (iEEG) study in humans found that co-firing of neurons in the Medial Temporal Lobe (MTL) at short delays predicted successful memory formation, whereas co-firing at longer delays predicted memory failure, which is consistent with STDP (Roux et al., 2021). Near-synchronous firing of hippocampal neurons should lead to effective synaptic connectivity, thus supporting the formation of associations between unrelated elements. Consistent with this prediction, studies in rodents have demonstrated that 40 Hz sensory stimulation using auditory tones or visual flicker can reach the hippocampus, as well as improving the hippocampal function, in turn improving cognitive function (Adaikkan et al., 2019; Martorell et al., 2019). We therefore speculate that neuronal changes in the hippocampus may be responsible for the STDP-like memory effect shown in the current study. Alternatively, the stimulus-timing dependent effect might be induced by multisensory integration that happens in primary sensory regions or higher-level association regions such as superior temporal sulcus (STS) or prefrontal cortex (Werner & Noppeney, 2010) which then have a knock-on effect on regions down-stream. This would be consistent with findings showing STDP to be ubiquitous in multisensory or large-scale cortical regions (Basura et al., 2015; Casula et al., 2016; Marks et al., 2018). Future experiments with good temporal and spatial resolution such as iEEG or MEG may reveal the underlying network of the stimulus-timing dependent memory effect we observe.

Our study suggests that, compared to the 20 ms stimulus peak interval, memory performance is significantly better when the stimulus peak interval is 6.67 ms. Future studies can test if the 20 ms interval is the boundary condition as the current study was limited by the modulation frequency, and therefore could not have an interval longer than 20 ms. It would be ideal if future studies can use a lower modulation frequency that can modulate the phase offset conditions to yield ∼ 7 ms, 20 ms, and longer than 40 ms peak intervals. Such a study could reveal if memory performance in the 40 ms interval condition is the baseline level, which means that the memory performance might be even worse than in the 20 ms interval condition, as predicted by the STDP rule. Furthermore, future experiments could pharmacologically manipulate the level of N-methyl-D-aspartate (NMDA) receptor activation (e.g. Weise et al., 2016), since STDP is NMDA dependent (Caporale & Dan, 2008; Mansvelder et al., 2019). If the stimulus-timing-dependent memory effect found in the current study is attributed to STDP, the difference between the memory performance in the shortest delay and longest delay conditions should be decreased, that is, the curves should be flattened when NMDA blockers are applied.

In conclusion, our study shows that sensory brain regions can be stimulated non-invasively with temporal offsets on the order of tens of milliseconds using rhythmic sensory stimulation to study human associative memory. The present results suggest a role of STDP for human episodic memory, which could be further investigated using iEEG or MEG approaches to reveal the underlying brain networks, as well as neuropharmacological studies to determine the underlying cellular mechanisms. Equally important, our study offers a precise and practical method to study the behavioural consequences of synaptic changes in human brain, which bridges the gap between the *in-vitro* studies and human episodic memory.

## Methods

### Participants

In total, 107 healthy English-speaking young adults participated in the experiments. In the auditory cue group, 51 participants (35 females; mean age: 19.6 years; range: 18 – 32 years) performed the experiment. Six participants were left-handed. The remaining 45 participants were right-handed. 34 participants were given course credits via the University of Birmingham’s Psychology Research Participation Scheme. The remaining participants were paid £8 per hour for their participation. The data from three participants were excluded because of chance-level memory performance. The data from the remaining 48 participants were retained for the final data analysis. In the visual cue group, 55 participants (35 females; mean age: 25 years; range: 18 – 40 years) participated in the experiment. 51 participants had not participated in the auditory cue group. Four participants who participated in the auditory cue group took part in the experiment as the study was originally designed as a within-group design. However, because of the COVID-19 pandemic, participants were not able to return. Four participants were left-handed. One participant was ambidextrous. 50 participants were right-handed. Apart from seven participants who were granted course credits, the remaining 48 participants were paid £8 per hour for their participation. The data from seven participants were excluded due to chance-level memory performance. The data from the remaining 48 participant were retained for the final data analysis. The EEG data from one participant were excluded due to poor EEG data quality (less than 15 trials were survived after artefacts rejections per condition). All participants had normal or corrected-to-normal vision and normal hearing.

### Materials

The visual stimuli were taken from the same stimulus set as those used in Clouter et al. (2017), Wang et al. (2018) and Chen et al. (2021). Some of the auditory stimuli were from the same set as those used in Clouter et al. (2017) and Wang et al. (2018). The remainder of the auditory stimuli were acquired from additional tracks of Apple Loops for Garage Band and two unique soundtracks that were royalty free. Movie clips of 3 s each had 227 frames in total with a frame frate of 75 frames/s. 288 movie clips were modulated at 37.5 Hz with luminance changing between 0 and 100% (initially starting at 100% luminance). All 288 sound clips were preprocessed using Audacity software (2.1.2 https://www.audacityteam.org/) as in Clouter et al. (2017). Each sound clip was presented concurrently with the presentation of a movie for 3 s, with a lag of 40 ms, which compensated for the fact that auditory stimuli are processed faster than visual stimuli Clouter et al. (2017). Sound amplitude was modulated at 37.5 Hz from 0 to 100% with a sine wave, at 0°, 90°, 180° and 270° phase offsets from the movies. The movie clips and sound clips were also modulated at 4 Hz. Each sound was modulated at 0° and 180° phase offsets from the four Hz sine-wave-modulated movies. Each sound clip was taken from one of the eight sound categories as described in Wang et al. (2018). Each sound category had 36 sound clips. All sounds were randomly divided into six sets of equal size (48 sounds per set), with the constraint that the number of sounds for each sound category was equal. The assignment of presentation frequency (37.5 Hz: 4 sets 192 sounds or 4 Hz: 2 sets 96 sounds) and phase offset conditions to each sound set were counterbalanced across participants. For each participant, a movie was assigned randomly at 37.5 Hz (192 movies), or 4 Hz (96 movies), then randomly assigned to a sound that was chosen to form a sound-movie presentation pair.

The experimental apparatus and stimulus presentation were identical to that used by Clouter et al. (2017) and Wang et al. (2018). The experiment was programmed with MATLAB (MathWorks) using the Psychophysics Toolbox extensions (Brainard, 1997; Kleiner et al., 2007; Pelli, 1997). Presentation of visual stimuli were on a 21-inch CRT display (Iiyama Vision Master Pro514 HM204DT) with an nVidia Quadro K620 graphics card (1058 MHz graphics clock, 2048 MB dedicated graphics memory, NVidia). Participants sat ∼60 cm from the centre of the monitor. 53 participants recorded with EEG while resting their head on a chin rest. The monitor screen refresh rate was 75 Hz. Auditory stimuli were presented with insert earphones (ER-3C, Etymotic Research). The physical presentation of phase offsets between movies and sounds, as well as the frequencies of the movies and sounds, were verified by a ThorLabs DET36A photodiode (https://www.thorlabs.com/) and a line-out speaker using 3.5 mm audio connectors connecting with a Biosemi Analogue Input Box (https://www.biosemi.com/aib.htm).

### Procedure

Participants in both the visual and auditory cue groups provided informed consent and were given task instructions, before practicing the procedure with four example trials. In the visual cue group, 29 participants were prepared for EEG data collection. The remainder of the participants in the visual cue group performed the behavioural tasks without EEG being recorded. In the auditory cue group, 24 participants’ EEG were recorded. 27 participants did the behavioural tasks without EEG recording. During the formal experiment, participants were monitored by a web camera connected to a monitor in the control room. Participants were asked to wave at the web camera if they had any questions or requests during the experiment.

The visual cue group were presented with 16 blocks of an associative memory task, where stimuli were modulated at 37.5 Hz. Each associative memory task block consisted of an encoding phase, a distractor phase, and an associative memory recall test phase. During the encoding phase, the procedure was the same as described in Wang et al. (2018). Participants were presented with a movie along with a sound for each trial. Each trial started with a fixation cross, which served as inter-trial-interval and lasted between 1 and 3 s. Then, the sound-movie pair was presented for 3 s. Participants were instructed to press one of the five number keys on a keyboard, to indicate how well the sound suited the contents of the movie after the presentation of the sound-movie pair. The instruction screen was presented until a response was made. The ratings ranged from 1 (the sound does not suit the contents of the movie at all) to 5 (the sound suits the movie very well). Participants were instructed to remember the association between the sound and the movie. Each block consisted of 12 trials. Four sounds from three categories were associated with the 12 movies with the constraint that the number of sounds for each phase offset condition was equal (i.e., one for 0°, one for 90°, one for 180° and one for 270° in one sound category). For participants whose EEG was recorded, another 8 blocks followed with only the encoding phase, during which the stimuli were modulated at 4 Hz. Participants were instructed to make a judgment as to how well the sound suited the contents of the movie but no memory test later on. Those blocks served as ground truth for the analysis of phase offsets between auditory and visual sources after adjusting for dipole orientation (see the section **Multimodal source reconstruction**).

The distractor phase was the same for both cue groups, as was done by Clouter et al. (2017) and Wang et al. (2018). During this phase, participants were presented with a random number that was drawn from 170 to 199 and instructed to count aloud backward from this number in steps of 3 for 30 seconds.

The associative memory recall test commenced following the distractor phase. In the visual cue group, the test phase consisted of 12 trials. Each trial started with a fixation cross between 1 and 3 seconds. Participants were presented with one of the 12 movies presented during the encoding phase for 3 s and instructed to recall the paired sound. Then, participants were presented with four sounds from the encoding phase and asked to select the sound that they would like to hear, using the number keys 1 through 4. After they completed listening to all four options, they were instructed to select the sound that they thought was played with the movie in the encoding phase, using the number keys 1 through 4. In both stages, the screen was presented until a response was made. The sounds from which to choose were all from the same sound category.

The associative memory task in the auditory cue group was similar to that of the visual cue group, except that each task block consisted of 16 trials and in total there were 12 associative memory task blocks. The stimuli in the 12 blocks were modulated at 37.5 Hz. The following six blocks consisted of the encoding phase only and the stimuli were modulated at 4 Hz if participants’ EEG was recorded. The different trial numbers between different cue groups were implemented because pilot results showed these trial numbers enabled participants to achieve acceptable memory performance and shortened the time required to complete the experiment. The encoding phase in each block of the auditory cue group was the same as in each block of the visual cue group. The associative memory recall test phase was identical to that employed by Wang et al. (2018). Participants were tested for all 16 trials within each block. Each trial started with a fixation cross randomly chosen to appear for between 1 and 3 s. Participants were presented with 1 of the 16 sounds presented during the encoding phase for 3 s, along with four still images from the four movies presented during the encoding phase. Participants were instructed to select the paired movie using the number keys 1 through 4. The instruction screen was presented until a response was made. The movies from which to choose were, in the encoding phase, all presented with a sound from the same sound category.

In both cue groups, two unimodal source localizer tasks were conducted following the multimodal blocks for those participants for whom EEG was recorded. The unimodal source localizer tasks served to separate sources from each sensory modality for multimodal source reconstruction. The tasks were exactly same as those used by Wang et al. (2018). The unimodal auditory task consisted of 50 trials of 4 Hz modulated sound clips. The unimodal visual task consisted of 50 trials of 4 Hz modulated movie clips. Participants were asked to rate how pleasant each sound or movie was using the number keys 1 (the sound or the movie was very unpleasant) through 5 (the sound or the movie was very pleasant) for each trial. The response to the 37.5 Hz stimulus was expected to originate from the same sensory source as the response from the 4 Hz stimulus. Only the unimodal localizers from the 4Hz modulated stimuli were used given that the signal-to-noise ratio for 4 Hz steady-state evoked responses was better than for 37.5 Hz responses.

For all participants, 24 sound-movie pairs randomly drawn from the associative memory task blocks were presented to test participants’ perception about the synchrony between a sound and a movie. The stimuli were modulated at 37.5 Hz. Participants were asked whether they could detect if the change of the modulated auditory stimulus was in synchrony with the corresponding modulated luminance of a movie (0° phase offset) or out-of-synchrony (90°, 180° or 270° phase offsets). Participants were instructed to press the number keys 1 for out-of-synchrony and 2 for in synchrony.

### EEG recordings and preprocessing

For participants whose EEG was recorded, 128 scalp channels of a BioSemi ActiveTwo system were used. Vertical eye movements were recorded from an additional electrode placed 1 cm below the left eye. Horizontal eye movements were recorded from two additional electrodes placed 1 cm to the left of the left eye and to the right of the right eye. Analogue signals of a photodiode that was attached to a white square informing the onset of a movie and two audio output channels of a sound were recorded by the BioSemi Analog Input Box, which resulted in three analogue signal channels (one for visual stimuli, two for auditory stimuli). These signals were recorded to later to correct for the onset of an EEG trigger, thus redefining the onset of an epoch by its actual onset. This is crucial for steady-state responses in higher frequency as a jitter of onset times might cause inaccurate phase and amplitude information of the fast evoked responses. Online signals were sampled at 2048 Hz using the BioSemi ActiView software. The position of each participant’s electrodes was tracked using a Polhemus FASTRAK device (Colchester) and recorded by Brainstorm (Tadel et al., 2011) implemented in MATLAB.

EEG data were preprocessed with the Fieldtrip toolbox (Oostenveld et al., 2011). The raw data less than 5 Gb were bandpass filtered between 1 and 120 Hz and bandstop filtered between 48-52 and 98-102 Hz to remove potential line noise at 50 and 100 Hz, then epoched from 2000 ms before stimulus onset to 5000 ms after stimulus onset. The raw data that were larger than 5 Gb were epoched first, then bandpass and bandstop filtered. The epoched data were downsampled to 512 Hz. An independent component analysis (ICA) was implemented after coarse artefact rejection of bad channels and trials by visual inspection. In the multimodal condition of the visual cue group, bad channels were excluded in five participants, with the average number of excluded channels being 4.2 (range 1 to 9). In the unimodal condition, bad channels were excluded in two participants (mean, 2.5, range 2 to 3). In the multimodal condition of the auditory cue group, bad channels were excluded in 10 participants (mean, 2.5, range 1 to 7). In the unimodal condition, bad channels were excluded in six participants (mean, 3.3, range 1 to 9). ICA components that indicated eye blinks and horizontal eye movements and regular pulse artefacts were removed from the data. The bad channels were interpolated by the method of triangulation of nearest neighbours based on the individuals’ electrode positions. After average re-referencing, trials that had artefacts were manually rejected by visual inspection. Participants with less than 15 artefact-free trials in any of the conditions of interest were excluded from further analysis. The mean numbers of trials remaining in each condition of the visual cue group and auditory cue group were listed in Table 1.

**Table 1.** Mean numbers of trials remaining in each condition in each cue group (values inside of brackets indicate range).

|  | Visual cue group | Auditory cue group |
| --- | --- | --- |
| Unimodal sound | 38 (22-49) | 38 (21-48) |
| Unimodal movie | 38 (23-50) | 38 (21-48) |
| Multimodal 37.5 Hz 0° | 35 (22-46) | 37 (26-45) |
| Multimodal 37.5 Hz 90° | 34 (22-44) | 38 (27-46) |
| Multimodal 37.5 Hz 180° | 36 (19-46) | 37 (21-44) |
| Multimodal 37.5 Hz 270° | 35 (23-44) | 38 (20-46) |
| Multimodal 4 Hz 0° | 38 (24-47) | 39 (28-46) |
| Multimodal 4 Hz 0° | 38 (19-47) | 38 (20-47) |

Since no individual MRI T1 structural scans were available, individuals’ electrode positions were aligned to a template head model. Then, source models were prepared with a template volume conduction model and the aligned individuals’ electrode positions.

### Unimodal source localisation

The unimodal source localisation was implemented in the same manner as described by Wang et al. (2018). The EEG data in the unimodal sound condition were transformed to scalp current density (SCD) using the finite-difference method (Huiskamp, 1991; Oostendorp & van Oosterom, 1996). The leadfields were also SCD transformed by applying the transformation matrix that was used for the SCD transformation (Murzin et al., 2013). Source activity was reconstructed using a linearly constrained minimum variance (LCMV) beamforming method (Van Veen et al., 1997). Time series SCD data were reconstructed in 2020 virtual electrodes for each participant. Source analysis was conducted with the SCD-transformed leadfields on the SCD-transformed data. In the unimodal movie condition, source analysis was implemented with leadfields that were computed based on scalp potentials. Time series potentials data were reconstructed in virtual electrodes for each participant. Event-related potential (ERP) was calculated at each virtual electrode for each unimodal condition. Time-frequency analysis was applied to the ERPs with a Morlet wavelet (width=7). Evoked power was averaged between 0.75 and 2.75 s after stimulus onset and between 3.5 and 4.5 Hz. A baseline condition was generated by randomly assigning each trial to 0°, 90°, 180° or 270° phase offset by moving the signal onset forward in time by 0, 32, 64 or 96 samples, which correspond to 0, 62.5, 125 or 187.5 ms with the constraint that the numbers of trials in each phase offset were approximately equal. The evoked power in the baseline condition was averaged between 0.75 and 2.75 s and between 3.5 and 4.5 Hz. The evoked power in each unimodal condition was normalised by subtracting the baseline evoked power from the condition evoked power and then divided by the baseline evoked power. This normalised evoked power was grand averaged across 48 participants of both visual cue and auditory cue groups. The grand average evoked power was interpolated to the MNI MRI template. The coordinates for the auditory and visual ROIs were determined by the locations of the maximum grand average evoked power.

### Multimodal source reconstruction

The multimodal source reconstruction was implemented as described by Wang et al. (2018) except for the steps of flipping the sign (i.e. adjusting the orientation of the dipoles) of reconstructed time series data. The multimodal data were SCD-transformed to reconstruct the time series data from auditory source to get cleaner time series source data from beamforming highly correlated sources (Murzin et al., 2013; Wang et al., 2018). Two sets of spatial filters based on the scalp electrodes over the right and left hemisphere, respectively, were computed. The SCD-transformed time series data was applied to the two sets of spatial filters and was extracted at the left auditory ROI and right auditory ROI which were predefined from the unimodal source localisation results. The time series data at the visual ROI was reconstructed without SCD transformation and was extracted at the visual ROI.

To solve the sign ambiguity (i.e. dipole orientation) problem caused by beamforming source reconstruction, the signs of reconstructed time series from each source were manually adjusted. The steps were as follows: first, the spatial filters that were extracted from three sensory ROIs were plotted on scalp. The spatial weights showed a preferred scalp distribution of corresponding sensory stimulation. Then, the signs of time series from the ROIs were set to be same as where the largest weights were distributed on the scalp, e.g., the time series data from the visual source would be applied by -1 if the occipital areas where the spatial weights were maximum showed a distribution of negative values. The sign would not be changed if the values of where the spatial weights were maximum were positive. The procedure was applied to all time series data regardless of experimental condition. The time series data from the auditory sources of the two hemispheres were averaged. A sanity check was performed using the data of the 4 Hz modulated conditions. The results were consistent with findings from Clouter et al. (2017) and Wang et al. (2018) in showing that the instantaneous phase difference in the 0° phase offset condition showed a preferred direction of 0° whereas the 180° condition showed a preferred direction of 180°. A control analysis was also performed to the data of 37.5 Hz modulated conditions to show that the phase relationships between 0° phase offset condition and other phase offset conditions in the auditory ROI were concentrated at 90°, 180°, and 270°. The phase differences between 0° phase offset condition and other phase offset conditions in the visual ROI always showed preferences at 0°. These results were consistent as the physical modulation of the sensory stimuli and remained constant between before and after the sign-flipping procedure.

Therefore, the flipping procedure did not bias the results in the direction of our hypothesis.

### Realigning condition labels

To compute the instantaneous phase differences, the source reconstructed time series data were bandpass filtered between 35 and 40 Hz. The stimulus onset (0 time point) was then redefined by the actual stimulus onset timing measured by the photodiode. The redefined epochs were cut by 1 s before stimulus onset and 4 s after stimulus onset. The ERPs were computed for each phase offset condition at each source. The Hilbert transformation was applied to the source grand averaged ERPs. The instantaneous phase differences were calculated between the unwrapped instantaneous phases from visual and auditory sources for 2 s, beginning 0.5 s after stimulus onset and ending 2.5 s after stimulus onset to avoid influences of onset and offset responses at gamma frequency. Rayleigh and V tests were used to test circular uniformity of the instantaneous phase differences in each phase offset condition. The condition labels were then realigned according to the actual instantaneous phase differences that each condition showed. The circular plots and statistics were done using the Circular Statistics Toolbox for Matlab (Berens, 2009). The ITPC analysis was done using the Fieldtrip toolbox. Time-frequency analysis was applied for each epoch in the realigned phase conditions at each source using multitaper time-frequency transformation based on Slepian sequences as tapers. The width of smoothing frequency was 1 Hz. The length of a sliding time-window was 1 s. The complex Fourier-spectra was computed using these parameters for frequencies of interest between 20 and 55 Hz and time of interest between -1 and 4 s. The complex Fourier-spectra was then normalised by its magnitude. The ITPC was computed by taking the magnitude of the mean of the normalised complex Fourier-spectra across trials. The ITPC for each condition at each source was grand averaged across 48 participants in both the visual cue and auditory cue groups. To compare the ITPC in each condition between each cue group at each source, the ITPC for each participant was averaged between 0.5 and 2.5 s and between 37 and 38 Hz.

On the single trial level, the instantaneous phase differences were calculated between the instantaneous phases from visual and auditory sources for the same length. The mean angle direction was computed across these data points. The angle values were sorted from -pi to pi and then divided into four equal sized bins (i.e. each bin containing the exact same amount of trials). If there were remainders after dividing the trial numbers by four, the same number of the last few trials whose angle values were closest to pi were discarded from the bin that had the largest angle values. Mean trial number in each bin for 48 participants is 36 (range from 23 to 45). Mean of the remainder trials is 1.6 (range from 0 to 3). The proportion of remembered trials in each phase bin was calculated. The ERPs were computed for each phase bin at each source. To check whether each phase bin showed preferred phase angle directions at 0°, 90°, 180° and 270°, the same procedure for computing the instantaneous phase differences between grand averaged ERPs at each source as described above was applied.

### Computational model

We adapted a computational neural network model from Wang et al. (2021), which comprises two groups of neurons that represent the neo-cortex (NC) and the hippocampus. Each group of neurons was split into two subgroups that represent the visual and auditory stimulus, respectively (N_nc_= 20, N_hipp_=10). The neuron physiology is simulated as in Wang et al. (2021) except that only the STDP learning rule was retained. There was no hippocampal theta learning system. Specifically, neuron membrane potential changes are simulated using an integrate-and-fire equation (Equation 1), where the membrane potential decays over time to a resting potential (E_L_ = -70mV) at a rate dictated by the membrane conductance (g_m_ = 0.03). Here, a spike event is generated if the voltage exceeds a threshold (V_th_ = -55mV), at which time the voltage is clamped to the resting potential for an absolute length of time to approximate a refractory period (2ms). As well as the leak current, the input current for model neurons contains the sum of all spike events occurring at pre-synaptic neurons (I_syn_), alternating current (AC) that represents NC alpha oscillations (I_AC_), any existing direct current (I_DC_) and an after-depolarisation (ADP) function (I_ADP_), described subsequently.

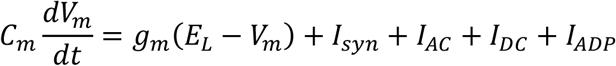

Equation 1 – The leaky integrate and fire equation.

Equation 2 explains the process by which neurons communicate through spike events, whereby the sum of all spike events over time makes up the I_syn_ current. Here, an alpha function is used to model the excitatory post-synaptic potential (EPSP), which provides an additive exponential function that diminishes the further the current time point (*t*) is from the initiating spike event (t_fire_). The amplitude of the function is dictated by the current synaptic weight of the post-synaptic synapse (0≤ *ρ* ≤1) multiplied by its maximal weight (W_max_). All spike events had a delay of 2 ms before they reached post-synaptic connections.

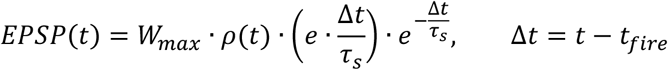

Equation 2 – Generation of an excitatory post-synaptic potential (EPSP) through time using an alpha-function.

Hippocampal neurons received additional input from an ADP function, as in previous models (Jensen et al., 1996; Parish et al., 2018); Equation 3; A_ADP_ = 0.2nA, τ_ADP_ = 250ms). This provided exponentially ramping input, which was reset after each spike-event (t_fire_).

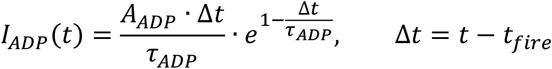

Equation 3 – After-depolarisation (ADP) function.

The learning rule was implemented via an adapted spike-time-dependent plasticity (STDP) mechanism, inspired by other models (Graupner & Brunel, 2012; Song et al., 2000). We consider two bi-directionally connected neurons in a traditional STDP framework. Upon the occurrence of a spike event in a model neuron, post-synaptic weights are strengthened for any given pre-synaptic neuron that spiked beforehand or weakened in the vice versa condition; the assumption being that the spike arriving at the post-synaptic connection must have either contributed to or competed with the spike event in question, depending on the directionality of the connection, leading to a reward or punishment of the synapse, respectively. To implement this, we calculate potential synaptic plasticity via functions for long-term potentiation (F_LTP_) and long-term depression (F_LTD_) at the time of an eliciting spike (t) in Equations 4.1-4.2.

In the case of potentiation (Equation 4.1), potential LTP at the post-synaptic connection (i) is calculated as the summation of historic pre-synaptic spikes (n_pre_) that occurred before the spike event in question (where t_i_ < t), weighted by an absolute value (A_+_ = 0.65). Contributions of pre-synaptic spikes were proportional to an exponential decay, thus favouring spikes that occurred close together in time (τ_s_ = 20ms). In the case of depression (Equation 4.2), potential LTD at the pre-synaptic connection (j) was similarly calculated as the summation of historic post-synaptic spikes (n_post_) that occurred before the spike event in question (where t_j_ < t), weighted by an absolute value (A_-_ = 0.65).

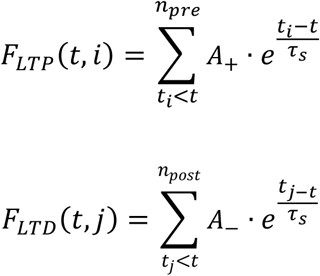

Equations 4.1 - 4.2 – Synaptic plasticity functions (F) calculate potential plasticity as the summation of the total number (n_post_ & n_pre_) of historic spike events (t_i_ & t_j_) arriving at a post- (i) or pre-synaptic (j) synapse relative to a given spike event (t), where an absolute value (A_+_/A_-_) is modulated by the difference in spike times and theta phase

Neurons within each subgroup (i.e., auditory, or visual) of the NC had a 25% chance of being connected (W_max_ = 0.3). Connections of neurons between subgroups were not implemented in NC as it was assumed visual and auditory stimuli had not been previously associated. Synaptic plasticity was also considered not to be operating on cortical synapses as in the complimentary systems framework (O’Reilly et al., 2014) it is assumed that cortical plasticity occurs on a much slower timescale. Background noise for each NC neuron was estimated by Poisson distributed spike-trains (4000 spikes/s, W_max_ = 0.023). A cosine wave of 10 Hz (amplitude = 0.1pA) was fed into NC neurons via I_AC_. Two constant inputs were fed into each NC subgroup to simulate presentation of visual and auditory stimuli via I_DC_ (amplitude = 1.75pA). These inputs were modulated by a cosine wave at 37.5 Hz with four phase offsets.

The two subgroups of hippocampal neurons that represented visual and auditory stimuli were fully connected to their NC counterparts (W_max_ = 0.35 for NC→Hip & W_max_ = 0.08 for Hip→NC synapses), as it was assumed both stimuli were previously known. Background noise for each hippocampal neuron was estimated by Poisson distributed spike-trains (1500 spikes/s, W_max_ = 0.015). Synapses within the entire hippocampus had a probability of 50% of forming a connection (W_max_ = 0.65), such that weights for intra-subgroup synapses were set to maximum and those for inter-subgroup synapses were initially set to 0. Synaptic plasticity was in effect on all hippocampal synapses, allowing for the association of visual and auditory stimuli to take place within the hippocampus.

Two cosine waves (−1≤ amplitude ≤1pA) were fed into the visual and auditory NC subgroups. The stimulus presentation length was three seconds (3000 data points). A 2-second inter-stimulus interval was used before visual-auditory stimulus presentation. The two cosine waves were modulated at 37.5 Hz with auditory stimulus phase offsets of 0°, 90°, 180°, and 270° from the visual stimulus. The simulations were run for 48 trials for each condition, which were then averaged across trials for each condition. For each simulation, we randomised a new set of initial synaptic connections as well as new Poisson distributed spike-trains for all conditions. To evaluate the recall performance of the model, the hippocampal weights after learning were averaged between 2.75 to 3 seconds after stimulus onset.

## Data availability

The datasets and the Matlab code generated during and/or analysed during the current study and the Matlab code of the model to run the simulations are available from: https://osf.io/fpyqk/.

## Supplementary Figures

**Figure S1.**
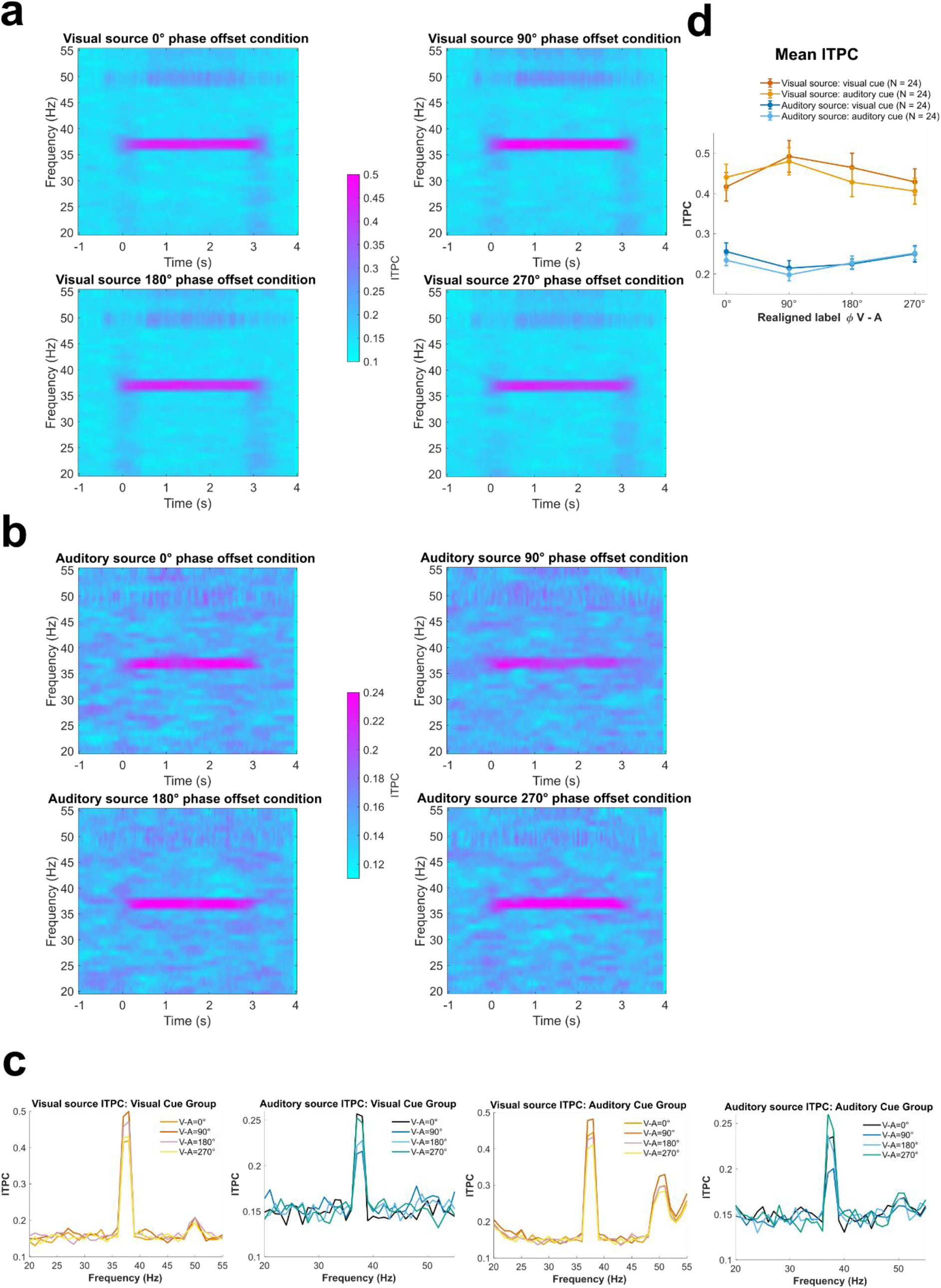
ITPC at each source during encoding phase. **(a)** Time frequency representations of ITPC averaged across 48 participants for each realigned phase offset condition at the visual source. **(b)** same as **(a)**but at the auditory source. **(c)** Spectrogram of ITPC averaged between 0.5 and 2.5 s of each cue group (N = 24 for both groups) for each realigned phase offset condition at each source. (**d**) Mean ITPC that was averaged between 0.5 and 2.5 s and between 37 and 38 Hz for each cue group (N = 24 for both groups) and each realigned phase offset condition at each source. Error bars represent SE. A significant interaction was found between brain region (visual vs. auditory) and realigned phase offset condition (90° vs. 270°), *F*(1,46) = 19.719, *p* < 0.001, η^2^_*p*_ = 0.3. In the visual cortex, ITPC in the 90° phase offset condition in which the visual stimulus led, was significantly stronger than in the 270° phase offset condition, *t*(47) = 3.758, *p* < 0.001 (one-tailed), Cohen’s *d* = 0.542. Whereas in the auditory cortex, ITPC in the 90° phase offset condition was significantly weaker than in the 270° phase offset condition, *t*(47) = -2.735, *p* = 0.004 (one-tailed), Cohen’s *d* = -0.395. There was no interaction between brain region, phase offset condition and cue group, *F*(1, 46) = 0.324, *p* = 0.572.

**Figure S2.**
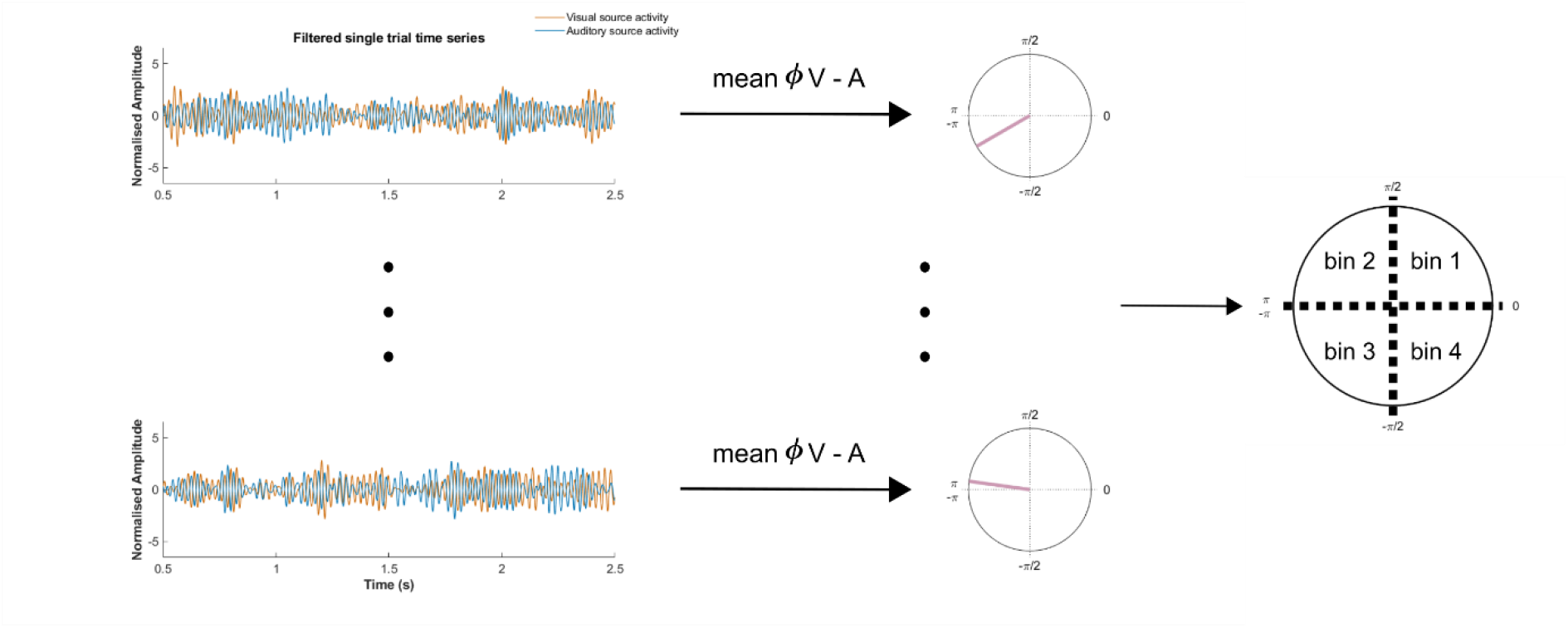
Procedure of single trial analysis. An example of sorting single trials into phase bins according to the instantaneous phase difference between band-pass filtered (35 – 40 Hz) visual and auditory EEG activity. The instantaneous phase differences were averaged between 0.5 and 2.5 s. All these values were sorted and divided into four equal phase bins. The trials in each bin should fall into the four quadrants of a cycle.

**Figure S3.**
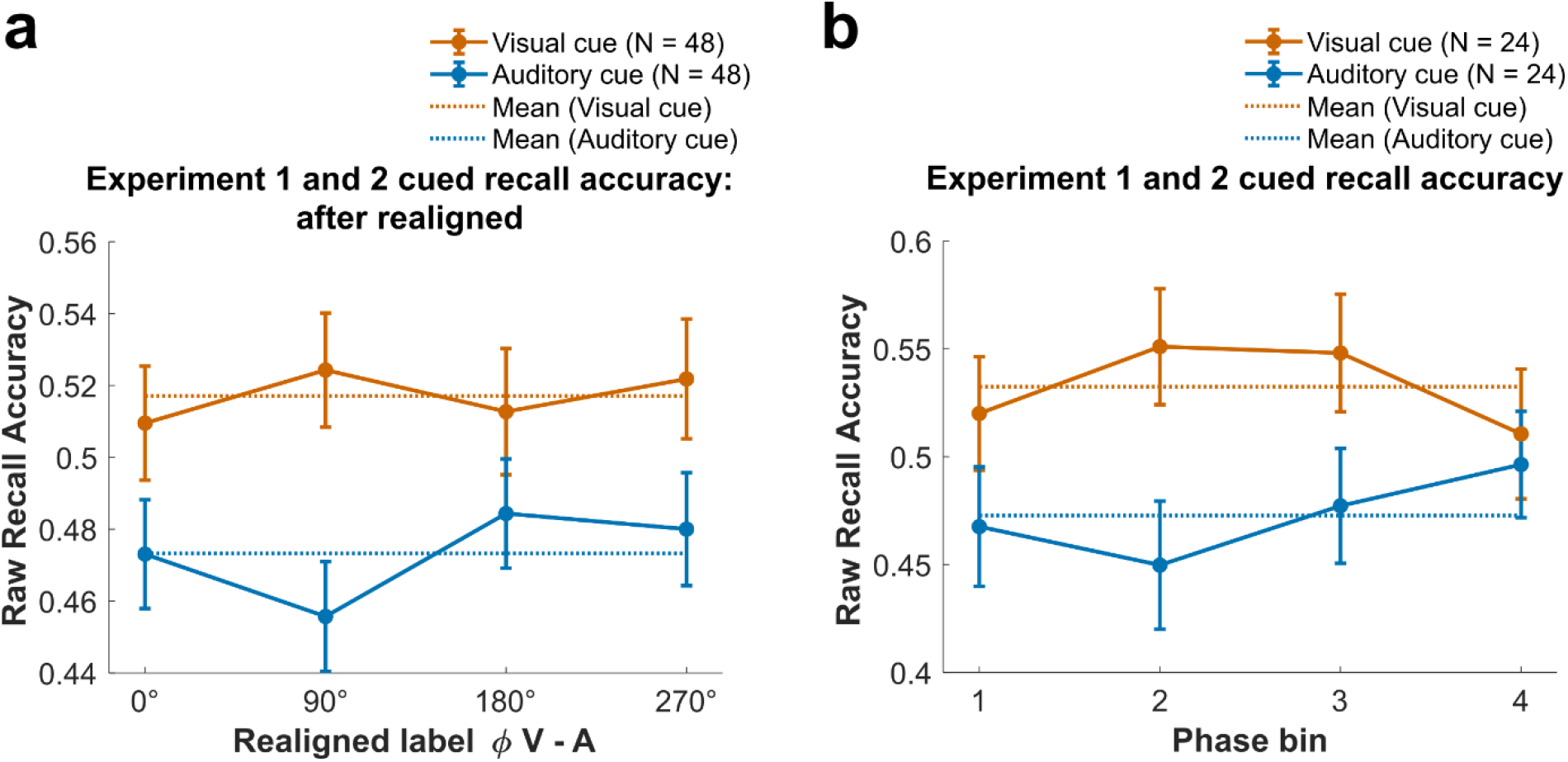
Behavioural results of raw recall accuracy. **(a)** Raw recall accuracy in each phase offset condition that showed the actual phase differences 0°, 90°, 180°, and 270° between visual and auditory grand average ERP, respectively. N = 48 for both groups. **(b)** Raw recall accuracy in each phase bin that was defined in the single trial sorting analysis. N = 24 for both groups. Error bars represent SE.

